# Biophysical properties and phenotypes of cell clusters detached from *Staphylococcus epidermidis* biofilms after matrix-targeted disruption

**DOI:** 10.64898/2026.01.28.701379

**Authors:** Sydney R. Packard, Gabriel Bulacan, Buddika Peiris, Randy Paffenroth, Elizabeth J. Stewart

## Abstract

Bacterial cells detached from *Staphylococcus epidermidis* biofilms are found to release predominantly as small oblate clusters (∼1.9 µm) in both untreated biofilms and biofilms treated with matrix-targeted disruptors. Quantitative image analysis common to colloidal science was applied to quantitatively evaluate the physical properties of 9,147 bacterial clusters detached from *S. epidermidis* biofilms with and without targeted disruption of individual matrix components (polysaccharides, proteins, extracellular DNA) or solubilization of the extracellular polymeric substances (EPS). Concentrations of *S. epidermidis* biofilm-detached cells are highest after matrix-targeted disruption of polysaccharides. *K-*means clustering, an unsupervised machine learning technique, was used to reveal that *S. epidermidis* biofilm-detached cells are released in five distinct phenotypes: small oblate, mid-sized oblate, large oblate, small spherical, and mid-sized prolate clusters. *S. epidermidis* biofilm detached cell clusters are predominantly oblate across three size groups (79.5%), with the small oblate phenotype representing 60.1% of cell clusters that have 3.1 ± 1.2 cells per cluster, Euclidean diameters of 1.9 ± 0.4 µm, anisotropy indices of 0.98 ± 0.05, and asphericities of −1.75 ± 0.31 on average. The proportion of *S. epidermidis* cell clusters within each biofilm-detached cell phenotype differs between matrix-targeted disruptors. There are also variations in the abundance of *S. epidermidis* biofilm detached cells after matrix-targeted disruption between growth conditions and strains. Evaluating the physical properties of biofilm-detached cells after matrix-targeted disruption is critical to understanding their translocation in fluid flow and susceptibility to the host immune response as well as in evaluating matrix-targeted disruption for biofilm control.

## Introduction

Bacterial biofilms are communities of surface-associated bacterial cells encased within self-produced matrix materials typically composed of polysaccharides, proteins, and extracellular DNA (eDNA)^1, 2^. The biofilm phenotype is prevalent across clinical, industrial, and natural environments^3^. Traditional techniques for removing biofilms range from mechanical removal^4^ (debridement^5^, pigging^6^, hydrodynamic shear^6^) to antimicrobial treatment^4^ (antibiotics^4^, chemical biocides^4, 6^, oxidants^4^). However, these biofilm removal strategies often fail to completely remove biofilms from surfaces leading to the resilience of biofilm communities^4, 6^.

Emerging methods for removing biofilms from surfaces focus on disrupting biofilm matrix materials^5, 7^. Matrix degradation enables biofilm disruption in hard-to-reach areas, such as biofilms in chronic wounds^5^ or pipes^7^. Matrix-targeted biofilm disruption can target individual components, such as polysaccharides, proteins, and extracellular DNA (eDNA)^5, 8^ or solubilization of the collective biofilm extracellular polymeric substances (EPS)^9^.

The efficacy of matrix-targeted biofilm disruption is frequently assessed by measuring the change in biofilm biomass after disruption using crystal violet (CV) staining^10, 11^ or optical density (OD) measurements of biofilms^12^ CV staining and OD measurements provide high-throughput evaluation of the relative effects of matrix disruption on biofilm biomass. However, these techniques do not provide quantitative data on properties of cells detached from biofilms after matrix disruption, which are critical for understanding bacterial survivability after matrix-targeted disruption.

Characterization of the properties of cell clusters detached from biofilms after matrix disruption is limited to enumerating the cell concentration with colony forming unit (CFU) counts^13–16^, evaluating released cell antimicrobial susceptibility^5, 15, 16^, and assessing released cell gene expression^15^. CFU counts provide estimates of surviving biofilm cells but often underestimate cell concentrations as one CFU can be cultured from either a single cell or larger cellular aggregate^17^. Antimicrobial susceptibilities of cells naturally dispersed from biofilms or released after biofilm disruption have been reported to be between those of planktonic cells and biofilms or similar to biofilms^15, 16^. Beyond viable cell counts, antimicrobial susceptibility, and cellular gene expression, the physical properties of bacterial cell clusters, such as size and morphology, play an important role in bacterial survivability and the efficacy of the host immune response^18, 19^.

Bacterial biofilms have been viewed as biocolloidal composite materials with rigid bacteria cells representing colloidal particles and matrix materials acting as a viscoelastic hydrogel^20^. Quantitative image analysis of bacterial biofilms using tools developed for characterization of colloidal systems^21^ and advanced image analysis algorithms^22^ has enabled evaluation of the cellular microstructure of bacterial biofilms^21, 22^. Similar quantitative image and particulate analysis techniques are used in colloidal science for characterizing cluster size and geometry (e.g. asphericity, anisotropy index) of self-assembled or aggregated clusters of colloidal particles^23, 24^, and are readily adaptable for evaluation of other colloidal systems, including characterization of bacterial cell clusters.

S*taphylococcus epidermidis* is one of the most common causes of biofilm infections on indwelling medical devices^25^. The *S. epidermidis* biofilm matrix is composed of polysaccharide intercellular adhesin (PIA), proteins, and extracellular DNA (eDNA)^26^. Chemical and enzymatic treatments such as sodium meta-periodate (NaIO_4_)^12, 27^, Proteinase K^12, 27^, DNase I^11, 28^ and high pH media^9^ have all been utilized for matrix-targeted disruption of *S. epidermidis* biofilms.

Here we use *S. epidermidis* biofilms as a model system to quantitatively evaluate physical properties, phenotypes, and antimicrobial susceptibilities of cell clusters released from biofilms after targeted disruption of individual matrix components (PIA, proteins, eDNA) and EPS solubilization. Biocolloidal properties of cells and cell clusters released from disrupted biofilms, including size and morphology, are assessed using confocal laser scanning microscopy coupled with quantitative image analysis. Cell clusters released from biofilms are then analyzed using *k*-means clustering, an unsupervised machine learning algorithm, to uncover hidden trends in the properties of detached cells. This work addresses two primary questions: (1) What is the impact of distinct matrix-targeted disruptors on the concentration, size, morphology, and antimicrobial susceptibility of bacterial clusters released from *S. epidermidis* biofilms? (2) What is the efficacy of matrix-targeted disruptors across *S. epidermidis* strains and growth environments? Evaluating the properties of cells detached from biofilms is a critical step toward understanding the survivability of biofilm cells and evaluating matrix-targeted disruption as a biofilm control strategy.

## Materials and Methods

### Bacterial strains and culture conditions

*S. epidermidis* RP62A (ATCC 35984), a PIA-dependent biofilm forming strain^29^ was used as the primary model organism. Additionally, *S. epidermidis* 1457 (ATCC 14990), also a PIA-dependent biofilm forming strain^29^, and six biofilm-forming *S. epidermidis* isolates (P2, P6, P12, P18, P37 and P47^30^, kindly provided by J.S. VanEpps, University of Michigan) were used. Strains were cultured using tryptic soy agar and tryptic soy broth supplemented with 1 wt. % glucose (TSB_G_).

### Biofilm growth conditions

*S. epidermidis* biofilms were grown in flow cells or 96-well plates. Flow cell biofilms were grown by seeding a flow cell with an overnight bacterial culture diluted to an OD_600_ of 0.1 (∼1 x 10^8^ cells/mL), followed by static incubation of the flow cell at 37°C for 45 minutes to allow for bacterial adhesion, and, finally, perfusion of the channel with TSB_G_ at a shear stress of 0.1 Pa for 24 hours. Flow cells had dimensions of 40 mm x 1 mm x 100 μm (length x width x height) and were fabricated with a polydimethylsiloxane (PDMS) channel bonded to a glass coverslip. A shear stress of 0.1 Pa represents the lower limit of the range of shear stresses in veins (0.1-1 Pa)^31^. For flow-cell grown biofilms, four experimental replicates were performed for each condition. Biofilms grown in polystyrene 96-well plates (CELLTREAT Scientific Products, Pepperell, MA), were seeded with an overnight culture of bacteria diluted to a concentration of ∼5 x 10^5^ cells/mL and incubated at 37°C and 60 RPM for 24 hours. For 96-well plate biofilm experiments, twelve experimental replicates were performed for each strain and treatment condition.

### Biofilm matrix disruption agents

Four previously established *S. epidermidis* biofilm matrix-targeted disruptors were used in this study. To disrupt polysaccharide intercellular adhesin (PIA), *S. epidermidis* biofilms were treated with 10 mM sodium meta-periodate (NaIO_4_) (Sigma-Aldrich, USA) in a 50mM sodium acetate oxidation buffer^12, 27^. Biofilm treatment with 50 mM sodium acetate alone does not significantly alter the abundance of cells detached from biofilms (Fig. S1). Biofilm matrix proteins were digested with 100 μg/mL Proteinase K (Invitrogen, USA) in TSB_G_ ^27, 28^. DNase I (Roche Diagnostics GmbH, Germany) in TSB_G_ (500 μg/mL)^28^ was used to disrupt matrix eDNA in *S. epidermidis* biofilms. Finally, pH 10 TSB_G_ was applied to solubilize the biofilm EPS, a method established for disassembling *S. epidermidis* biofilms by Anonymous et al.^9^. TSB_G_ alone was used for untreated biofilm experiments.

### Matrix-targeted disruption of S. epidermidis biofilms grown in flow cells

After 24 hours of growth, biofilms were treated for one hour with a matrix-targeted disruptor to induce targeted PIA, protein, or eDNA disruption or EPS solubilization. Flow cell effluent was collected in four, 15-minute intervals during the 1 hour treatment to limit effects of bacterial growth during effluent collection. *S. epidermidis* RP62A has a doubling time of 62 minutes at 37°C in TSB_g_ (Fig. S2).

### Confocal Laser Scanning Microscopy (CLSM) imaging and quantitative image analysis of biofilm biomass remaining in flow cells after treatment

The height and biomass of biofilms after matrix-targeted disruption was characterized using CLSM coupled with image analysis in COMSTAT – a Matlab program used to quantify biofilm structure^32^. Biofilms were stained with 5 μM of Syto40 (ThermoFisher Scientific™, USA S11351) for 30 minutes. Syto 40 has an excitation/emission of 420/441 nm. CLSM image volumes were acquired on a Leica Stellaris 8 CLSM with a 63x,1.40 NA oil immersion objective lens. Image voxels were 0.36 μm x 0.36 μm x 0.50 μm. For each experimental replicate, five image volumes were taken along the center of the flow cell with 1000 µm between images. Biofilm biomass is the number of pixels containing biomass in a 3D CLSM image volume multiplied by the voxel size and divided by the surface area of the image volume at the substrate (µm^3^/µm^2^)^32^. Biofilm height is reported as the maximum height containing pixels with biomass.

### Characterization of the concentration of cells detached from biofilms

For each of the four biofilm effluent collection timepoints, the concentration of cells released from biofilms was measured using a hemocytometer under a Zeiss Axiovert 200M fluorescence microscope with a 40x LD, 0.6NA, Ph2 objective lens, where the number of cells was counted in five 0.04mm^2^ hemocytometer squares.

### CLSM imaging and quantitative image analysis of cells detached from biofilms

The biofilm effluent collected at each timepoint was imaged in 8-well Nunc Lab-Tek II Chambered Coverglass dishes (ThermoFisher Scientific™, USA 155409). Effluent was stained with 1 μM Syto9 (ThermoFisher Scientific™, USA S34854) for 60 minutes and then imaged using a Leica Stellaris 8 confocal scanning laser microscope (CLSM) with a 100x, 1.44 NA, oil immersion objective lens. The excitation/emission wavelengths for Syto9 are 485nm/498nm. Image volumes were 40.96 μm x 40.96 μm in the x-y plane and between 5-20 μm in the z-direction (height), where image height was based on the maximum cell cluster height within the image volume. Image voxels were 80 nm x 80 nm x 80 nm. Biofilm-detached bacterial clusters collected after no treatment, NaIO_4_ treatment, and Proteinase K treatment were immobile on untreated coverglass. After DNase I treatment, biofilm detached cell clusters were immobile on coverglass treated with oxygen plasma (100 watts, 45 seconds). After pH 10 TSB_G_ treatment, biofilm detached cell clusters were immobile on coverglass treated with Rain-X Glass Water Repellent (Illinois Tool Works, Inc., USA). Coverglass treatment (untreated, oxygen plasma, or Rain-X Glass Water Repellent) altered the mobility of settled bacterial cell clusters (Fig. S3), but did not alter cluster size, Euclidean diameter, anisotropy index or asphericity of *S. epidermidis* RP62A planktonic cell clusters (Fig. S4). Five image volumes were taken at each of the four collection timepoints for a total of 20 image volumes per experimental replicate and 80 image volumes per experimental condition.

Quantitative image analysis of CLSM image volumes was utilized to identify the 3D spatial positions of cells within clusters and determine physical properties of cell clusters detached from biofilms. Identification of bacterial centroid locations within CLSM image volumes was performed using TrackPy^33^ – a Python image analysis package based on the Crocker and Grier algorithm for identifying colloidal particles^34^. Briefly, raw 3D image volumes (Fig. 1A) were preprocessed using a Gaussian mask to subtract background noise from images, candidate particle locations were identified as the brightest pixels within a region of a cell’s approximate radius, and particle locations were refined to the bacterium centroids in 3D (Fig. 1B)^34^. Using the methods of Savin and Doyle^35^, the static error in particle location was determined to be ± 23 nm in the object plane (xy) and ± 56 nm in the axial plane (xz) (Fig. S5).Following bacterial centroid identification, 3D spatial coordinates of bacterial centroids were grouped into clusters and analyzed using the freud library^24^. To quantify the number of cells per cluster, all cells within a given image volume were grouped according to the freud cluster module where cells are grouped into the same cluster if their centroids are within a specified radius of another cell (Fig. 1C)^24^. The cluster cutoff distance (*r_c_*) was chosen to be 1.5 μm based on the inflection point on the cluster distribution curve (Fig. S6).

**Figure 1.**
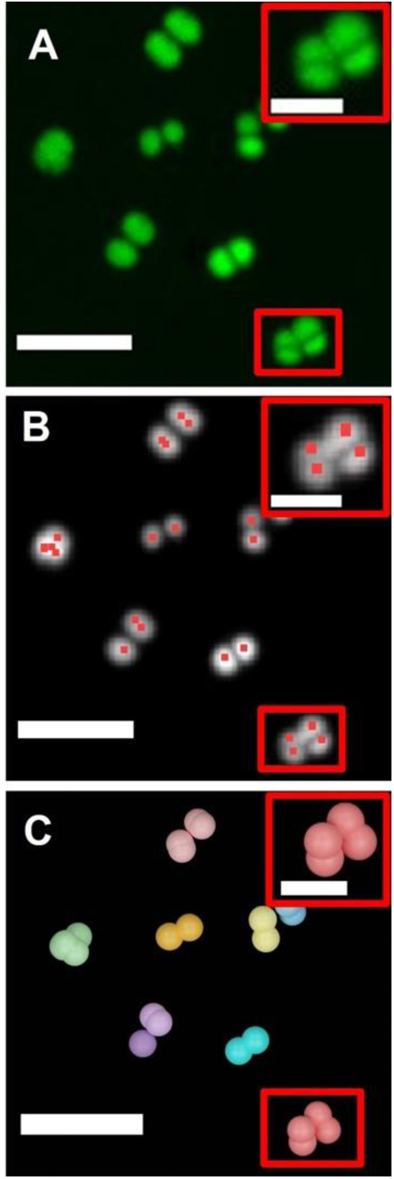
Identification of cell clusters detached from *S. epidermidis* biofilms using quantitative image analysis of confocal laser scanning microscopy (CLSM) images. Projection of maximum intensities of a representative (A) raw and (B) filtered CLSM image section (16.64 x 16.64 x 5.36 µm^3^) of bacterial cells detached from a biofilm. Bacterial centroids are identified by red squares. (C) Renderings of bacterial cells grouped into bacterial clusters with a cluster cutoff (r_c_) of 1.5 µm within the representative image volume, where bacteria in the same cluster are the same color. Bacteria centroids are rendered with a 1.2 µm sphere. Scale bars = 5 µm. Insets are of the cluster within the red box on the image. Scale bars = 2 µm.

The size of cellular clusters was calculated as the maximum Euclidean distance, *d*, between the two furthest cells, denoted as *p* and *q*, in each cluster (Eq. 1). To account for the full size of a bacterial cell, twice the radius of a single bacterial cell (*r* ∼ 350 nm) is added to the distance between *p* and *q*, as *p* and *q* are distances between bacterial centroids:

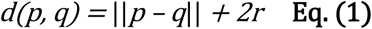

Asphericity (*b*) and shape anisotropy index (*κ^2^*) and were used to evaluate cellular cluster shape. Asphericity provides a quantitative metric describing whether a cluster of points are oblate (*b* < 0), spherical (*b* = 0), or prolate (*b* > 0) and shape anisotropy index provides a measure of whether an object is spherical (*κ^2^* = 0) or linear (*κ^2^*= 1). Cluster shape anisotropy index and asphericity were calculated in 3D from the radius of gyration, *R_g,_* in the freud Python library using the following equations^36^:

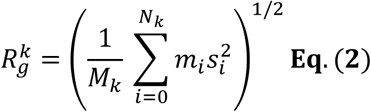

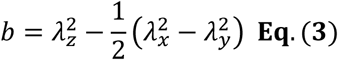

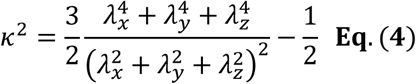

where *M_k_* is the total mass of the particles in the *k*^th^ cluster, *N_k_* is the number of particles in the *k*^th^ cluster, *m_i_* is the mass of particle *i*, *s_i_* is the distance of particle *i* from the center of mass of the *k*^th^ cluster, and λ_x_, λ_y_, and λ_z_ are square roots of eigenvalues of the gyration tensor from the radius of gyration.

### Classification of biofilm-detached cell clusters using k-means clustering analysis

The properties of *S. epidermidis* cell clusters detached from all untreated and treated biofilms were analyzed using *k*-means clustering^37^. *k*-means clustering partitions data into a user-defined number of groups, *k*, based on proximity to neighboring datapoints, thereby, sorting datapoints based on similarity to other datapoints^37^. Input ‘features’ labeled for each datapoint in the *k-*means clustering model were: number of cells per cluster, Euclidean diameter, anisotropy index, and asphericity. The dataset was scaled using a z-transform to standardize the weight of each ‘feature’ prior to performing *k*-means clustering. *k-*means clustering was performed with *k-*values from 1 to 12, plotted against the sum of square error, and the optimal *k-*value was identified using the Elbow method^37^. While *k-*means clustering is an unsupervised machine learning method for clustering data into user-defined groups, the stability of the identified *k-*value was assessed by performing *k-*means clustering on five random subsets of the data akin to 5-fold cross validation^38^. A stable *k-*value captures the inherent structure of the data and contains *k*-means groups that remain consistent across different subsets of data^39^. *k-*means clustering was performed on the full dataset and across five sub-sections of the data to evaluate stability of the *k*-value. The mean, standard error of the mean, and range of: number of cells per cluster, maximum Euclidean diameter, anisotropy index, and asphericity for each *k*-means group were calculated. t-distributed stochastic neighbor embedding (t-SNE), a data visualization technique for mapping statistically similar and dissimilar datapoints from high-dimensional data in two-dimensional space ^40^, was used to visualize phenotypes of bacterial cell clusters for the optimal *k-*value.

### Determination of the minimum inhibitory concentration (MIC) of vancomycin against biofilm detached cells

The MICs of vancomycin toward cells detached from *S. epidermidis* RP62A biofilms after targeted matrix disruption were determined using the CLSI standard^41^. Briefly, the effluent of *S. epidermidis* biofilms grown in flow cells was collected for 1 hour with and without treatment with biofilm matrix disruption agents. Released cell effluent was diluted to a final concentration of 5 x 10^5^ cells/mL and seeded with 0.5, 1.0, 2.0, 4.0, 8.0, 16.0, 32.0 or 64.0 µg/mL vancomycin in TSB_G_ in 96-well plates. After 24 hours, the MIC was determined by visual inspection as well as OD_600_ measurements.

### Matrix-targeted disruption of S. epidermidis biofilms grown in 96-well plates

To characterize the abundance of cells released from *S. epidermidis* RP62A, 1457, P2, P6, P12, P18, P37 and P47 biofilms after matrix-targeted disruption, the supernatant of biofilms grown in 96-well plates was aspirated and replaced with 200µL of TSB_G_, 10 mM NaIO_4_ in 50 mM sodium acetate buffer, 100 µg/mL Proteinase K in TSB_G_, 500 µg/mL DNase I in TSB_G_, pH 10 TSB_G_ or 50 mM sodium acetate buffer. Abundance of cells in 12 wells per treatment condition was evaluated using OD_600_ measurements of the biofilm supernatant with five technical replicates per biofilm supernatant sample. Relative abundance of cells released from biofilms after treatment was quantified as the ratio of treated to untreated biofilm released cell abundance.

### Statistical Analysis

Statistical analysis was performed using GraphPad Prism version 9 (San Diego, CA, USA). Data normality was assessed for all datasets using Shapiro-Wilk tests. For normally distributed data (concentration of cells detached from biofilms, remaining biofilm biomass, biofilm height), statistical significance was determined using ordinary one-way ANOVA followed by Dunnett’s T3 multiple comparisons tests to compare untreated biofilms against each treatment condition (NaIO_4_, Proteinase K, DNase I, pH 10 media). For datasets with non-Gaussian distributions (number of cells per cluster, Euclidean diameter, anisotropy index and asphericity of cell clusters detached from *S. epidermidis* biofilms across treatment conditions (Fig. S7) and planktonic cell clusters on untreated and treated coverglass (Fig. S4)), non-parametric Kruskal-Wallis tests with Dunn’s multiple comparison tests were employed. To compare untreated and treated biofilms between *S. epidermidis* strains (RP62A, 1457, P2, P6, P12, P18, P37, P47), two-way ANOVA with Dunnett’s T3 multiple comparison tests was used. For all statistical analyses, significance was defined as *P ≤ 0.05, **P ≤ 0.01, ***P ≤ 0.001, and ****P ≤ 0.0001.

## Results

Matrix-targeted disruption of *S. epidermidis* RP62A biofilms grown at a shear stress of 0.1 Pa results in the release of cells and cellular clusters with concentrations distinct from untreated biofilms, where disruption of PIA – the predominant matrix component of the biofilms – releases seven times more cells than in undisrupted biofilms, 2.8 x 10^8^ cells/mL compared to 4.1 x 10^7^ cells/ mL (Fig. 2A). Disrupting eDNA in the biofilm matrix and collective EPS solubilization of *S. epidermidis* biofilms has a more modest effect on released cell concentrations with 3.2 times and 3.1 times more biofilm cells released during treatment than in untreated biofilms, respectively (Fig. 2A). The concentration of cells released after protein-targeted disruption of *S. epidermidis* biofilms is comparable to the cell concentration released from untreated biofilms (Fig. 2A). The vancomycin MIC of biofilm released cells is the same as the MIC for planktonic *S. epidermidis* cells (2 µg/mL) after targeted disruption of proteins, eDNA, and total matrix solubilization (Table S1).

**Figure 2.**
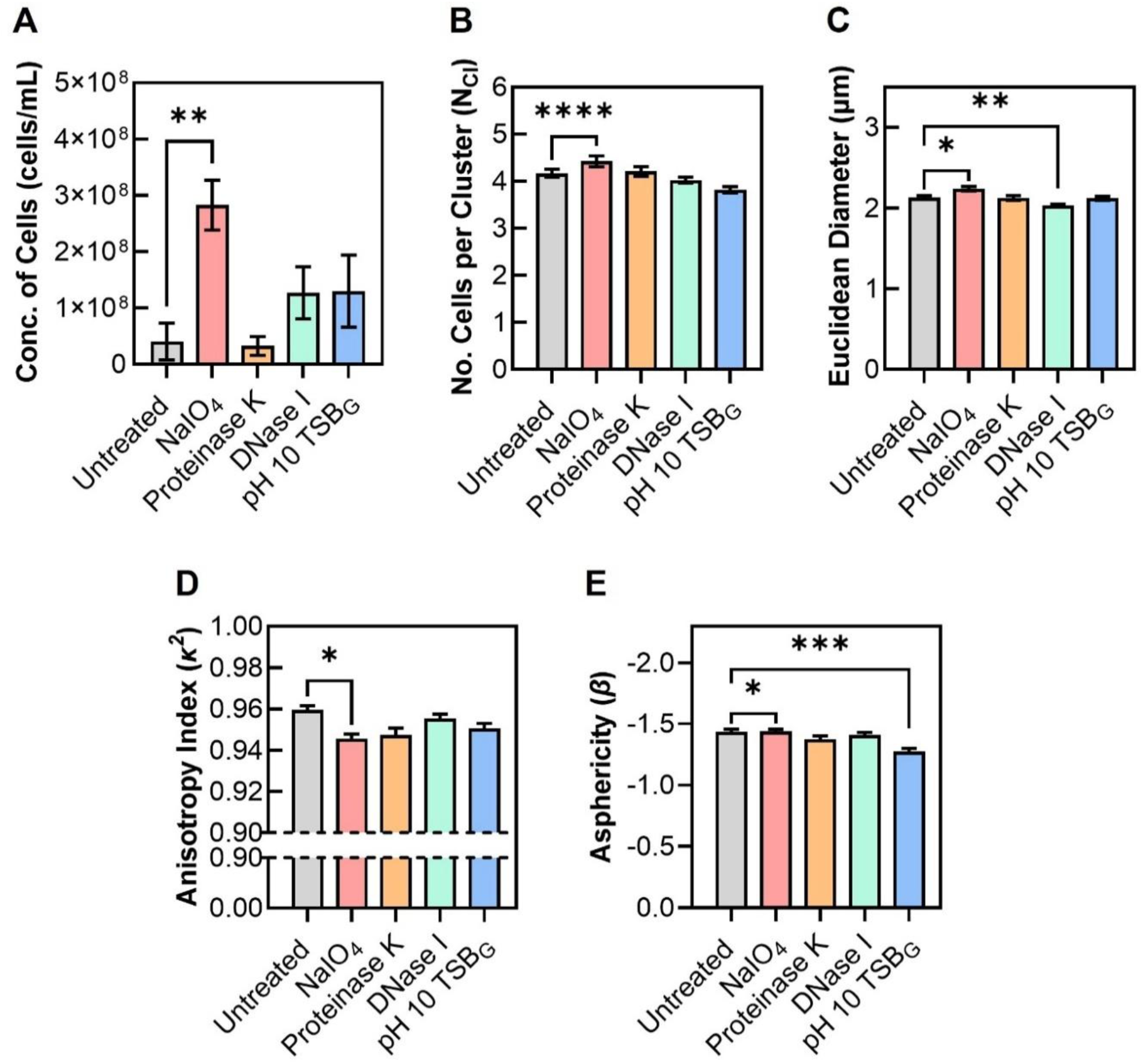
Average physical properties of cells and cell clusters detached from *S. epidermidis* biofilms after matrix-targeted biofilm disruption. (A) Average concentration of cells detached from biofilms. Statistical significance determined by one-way ANOVA with Dunnett’s T3 multiple comparisons tests (**P ≤ 0.01). (B-E) Average (B) number of cells per cluster, (C) Euclidian diameter, (D) anisotropy index, and (E) asphericity of cell clusters detached from *S. epidermidis* RP62A biofilms with no treatment or treated with NaIO_4_, Proteinase K, DNase I, or pH 10 TSB_G_ with statistical significance determined by non-parametric Kruskal-Wallis tests with Dunn’s multiple comparison tests (*P ≤ 0.05, **P ≤ 0.01, ***P ≤ 0.001, ****P ≤ 0.0001).

Changes in *S. epidermidis* RP62A biofilm structure due to matrix-targeted disruption are also most significant after PIA-targeted disruption, where biofilm biomass and height are 4.7 times and 1.6 times lower than the biomass and height of untreated biofilms, respectively (Fig. S8). This reduction in biofilm biomass and height aligns with the observed significant increase in cells released from biofilms after PIA-targeted biofilm disruption (Fig. 2A). The variable effects of different biofilm matrix-targeted treatments are less pronounced in the average biofilm biomass and height (Fig. S8) than in the concentrations of cells released from biofilms after protein-targeted disruption, eDNA-targeted disruption and EPS solubilization of *S. epidermidis* RP62A biofilms (Fig. 2A).

The average size and morphology of biofilm-detached cell clusters from both untreated and matrix-disrupted *S. epidermidis* RP62A biofilms are similar (Fig. 2B-E). Cell clusters detached from *S. epidermidis* biofilms contain 3.8 to 4.4 cells per cluster (Fig. 2B) and are between 2.0 – 2.3 µm in diameter on average (Fig. 2C) across treatments. *S. epidermidis* biofilm detached cell cluster morphologies are primarily linear and oblate with average anisotropy indices between 0.94 and 0.96 (Fig. 2D) and average asphericities between −1.44 and −1.28 (Fig. 2E) across all treatments. The most significant differences in the size and morphology of *S. epidermidis* RP62A cell clusters detached from biofilm are for PIA-targeted disruption, where cell clusters were collectively slightly larger (Fig. 2B, 2C, Fig. S7A), less linear (lower anisotropy index values, Fig. 2D), and more oblate (lower asphericity values, Fig. 2E) than cell clusters released from untreated *S. epidermidis* biofilms. There are also small, yet statistically significant, differences in the Euclidian diameters of cell clusters after DNAse I treatment (Fig. 2C) and the asphericities of cell clusters after pH 10 TSB_G_ treatment (Fig. 2E) as compared to untreated biofilms.

While average *S. epidermidis* biofilm released cell cluster sizes and morphologies are similar across matrix-targeted treatments, the range and distributions of the number of bacterial cells per cluster, cluster Euclidean diameters, anisotropy indices, and asphericities are broad and non-Gaussian (Fig. S7). Thus, *k-*means clustering, an unsupervised machine learning algorithm, is used to uncover hidden trends in *S. epidermidis* biofilm-detached cell cluster properties.

Five distinct *S. epidermidis* biofilm-detached cell cluster phenotypes, or *k-*means groups are revealed through *k-*means clustering analysis of all *S. epidermidis* RP62A cellular clusters detached from untreated and treated biofilms (n = 9,147 biofilm-detached cellular clusters). The five phenotypes are visualized in low-dimensional space using t-SNE visualization where phenotypes 1, 2, and 3 cluster closely together, and phenotypes 4 and 5 form distinct groups (Fig. 3A). This suggests that phenotypes 1, 2 and 3 share similar characteristics that vary gradually along a continuum, contrasting with the sharp categorical separation for phenotypes 4 and 5 (Fig. 3A). Five is the optimal number of *k*-means groups for the dataset as a *k-*value of 5 is the maximum point of curvature in a plot of the *k*-value against the normalized sum of squared error (Fig. S9A). A *k*-value of 5 also captures the inherent structure of the dataset as the average values of each feature (number of cells per cluster, Euclidean diameter, anisotropy index, and asphericity of the cellular clusters) are distinct across the 5 *k-*means groups and are consistent and stable across the 5-fold sub-samples of dataset (n = 1,829 biofilm-detached cellular clusters per sub-sample) (Fig. S9B-E).

**Figure 3.**
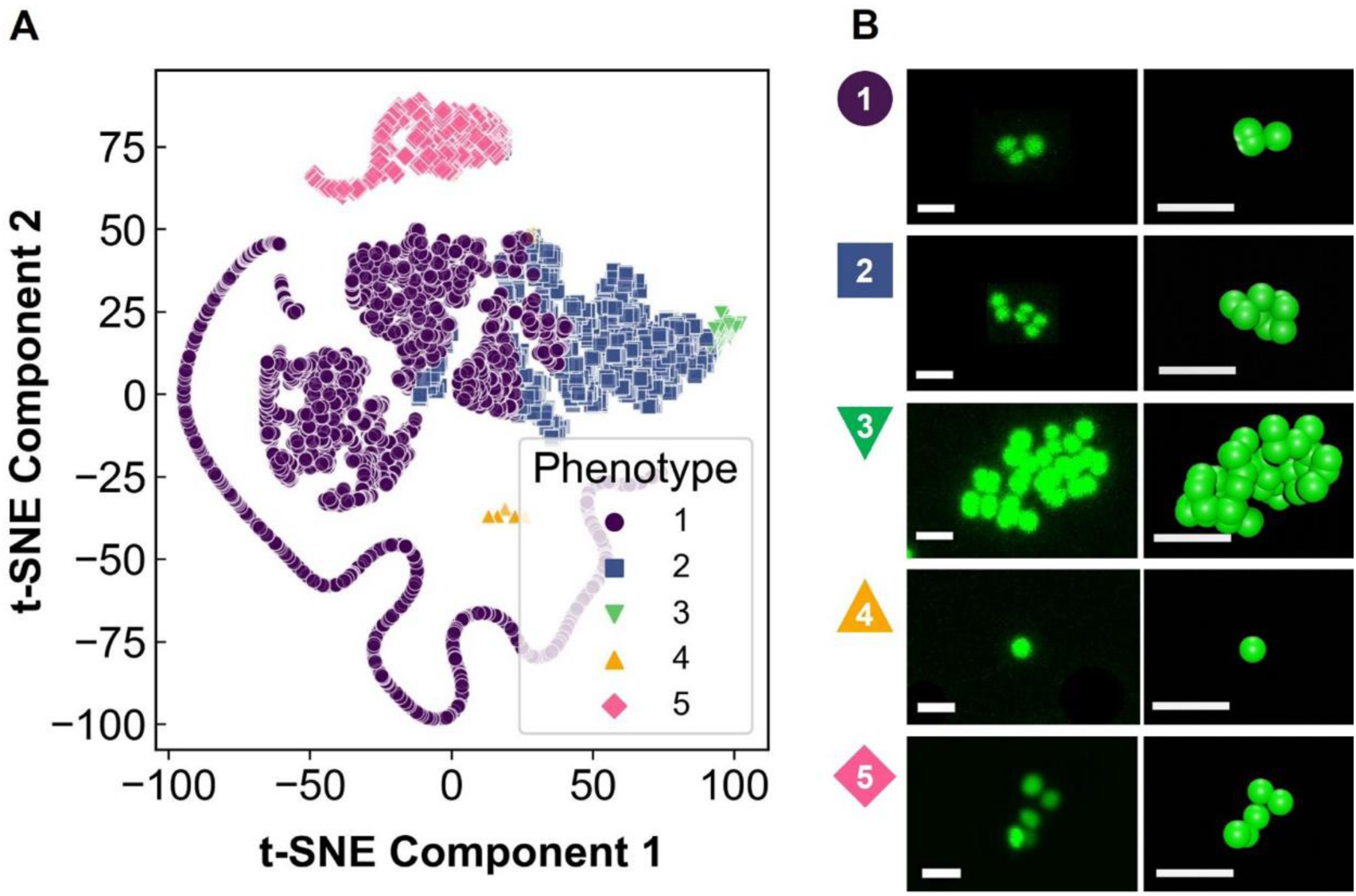
*k-*means clustering of the biophysical properties of cellular clusters detached from *S. epidermidis* RP62A biofilms. (A) Two-dimensional t-distributed stochastic neighbor embedding (t-SNE) visualization of the five distinct biofilm detached cell phenotypes. (B) Projections of 3D CLSM images (left column) and renderings (right column) of representative bacterial cell clusters for each biofilm-detached cell cluster phenotype. Scale bars = 2 µm. Phenotype 1 (purple circle) contains small, oblate clusters, Phenotype 2 (blue square) contains mid-sized, oblate clusters, Phenotype 3 (green inverse triangle) contains large, oblate clusters, Phenotype 4 (orange triangle) contains small, spherical clusters, and Phenotype 5 (pink diamond) contains small, prolate clusters.

The five *S. epidermidis* biofilm-detached cell cluster phenotypes have three distinct morphologies (oblate, prolate, spherical) and three size groups (small, mid-sized, large) (Fig. 3B, Table 1). The morphology of 79.8% of *S. epidermidis* biofilm-detached cell clusters is linear and oblate, where linear, oblate cell clusters are grouped into three distinct size groups: small, mid-sized, and large. The small, linear, oblate group (Phenotype 1) contains the highest percentage of clusters (60.4%), the mid-sized, linear, oblate group (Phenotype 2) contains the second highest percentage of clusters (18.1%), and the large, linear, oblate group (Phenotype 3) contains the lowest percentage of clusters (1.3%) (Table 1). For the small, mid-sized and large oblate cluster phenotypes, the average number of cells per cluster are 3.1 ± 1.2 cells, 7.5 ± 2.7 cells, and 27.9 ± 16.7 cells, and the average cluster diameters are 1.9 ± 0.4 µm, 3.4 ± 0.8 µm, and 6.9 ± 2.1 µm, respectively (Phenotypes 1, 2, 3; Table 1). As cluster size increases from small to large within the three linear, oblate phenotypes, cell clusters become less linear as indicated by anisotropy index and asphericity values moving toward zero (Table 1). The mid-sized and large linear, oblate cluster phenotypes (Phenotypes 2, 3) contain a small sub-set of prolate clusters as some clusters have asphericity values greater than zero (Table 1). The remaining 20.2% of *S. epidermidis* biofilm released cell clusters fall into two additional *k-*means phenotypes: small, spherical clusters (Phenotype 4, 11.3% of clusters) and mid-sized, prolate clusters (Phenotype 5, 8.9% of clusters) (Fig. 3B, Table 1). The small, spherical clusters (Phenotype 4) have average diameters of 0.7 ± 0.1 µm and consist of predominantly individual cells (99% (1030/1037) of clusters). The mid-sized, prolate clusters (Phenotype 5) have clusters with diameters of 5.6 ± 2.5 µm and 2.6 ± 0.7 cells/cluster on average (Table 1). Thus, *S. epidermidis* cells released from biofilms are predominantly linear and oblate in geometry (79.8%) or small (71.7%) in size with large clusters representing the smallest fraction of biofilm-detached cells (1.3%) (Table 1).

**Table 1.** Five phenotypes of *S. epidermidis* RP62A biofilm detached cell clusters identified using *k*-means clustering analysis.

| ***k*-Means Phenotype** |  |  | **No. of cells per cluster<sup>a</sup>** | **Euclidean Diameter<sup>a</sup> (μm)** | **Anisotropy Index<sup>a,b</sup>** | **Asphericity<sup>a,c</sup>** | **Percentage** |
| --- | --- | --- | --- | --- | --- | --- | --- |
| **No.** | **Size** | **Morphology** |  |  |  |  |  |
| 1 | Small | Linear, Oblate | 3.1 ± 1.2<br>(2 to 8) | 1.9 ± 0.4<br>(0.7 to 3.3) | 0.98 ± 0.05<br>(0.51 to 1.00) | -1.75 ± 0.31<br>(-2 to -0.55) | 60.4 % |
| 2 | Mid-sized | Linear, Oblate | 7.5 ± 2.7<br>(4 to 19) | 3.4 ± 0.8<br>(2.1 to 6.8) | 0.87 ± 0.11<br>(0.38 to 1.00) | -1.30 ± 0.33<br>(-1.98 to 0.73) | 18.1 % |
| 3 | Large | Linear, Oblate | 27.9 ± 16.7<br>(13 to 131) | 6.9 ± 2.1<br>(4.1 to 20.2) | 0.75 ± 0.13<br>(0.40-0.99) | -0.76 ± 0.73<br>(-1.89 to 0.86) | 1.3 % |
| 4 | Small | Spherical | 1.0 ± 0.4<br>(1 to 8) | 0.7 ± 0.1<br>(0.7 to 2.6) | 0.00 ± 0.02<br>(0.00 to 0.38) | 0.00 ± 0.03<br>(-0.46 to 0.32) | 11.3 % |
| 5 | Mid-sized | Linear, Prolate | 5.6 ± 2.5<br>(2 to 19) | 2.6 ± 0.7<br>(0.7 to 5.5) | 0.90 ± 0.12<br>(0.27 to 0.30) | 0.88 ± 0.15<br>(-0.02 to 1.0) | 8.9 % |

<sup>a</sup>Mean ± SEM (Range)
<sup>b</sup>Anisotropy index of 0 is spherical and 1 is linear.
<sup>c</sup>Asphericity > 0 is prolate, 0 is spherical, and < 0 is oblate.

The fraction of *S. epidermidis* biofilm-released cell clusters from each of the five phenotypes varies between untreated biofilms and biofilms treated with each matrix-targeted disruptor (Table 2). PIA-targeted biofilm disruption releases the highest percentage of large, linear, oblate clusters (Phenotype 3, 3.1%) and small, spherical clusters (Phenotype 4, 14.0%) as compared to the other biofilm matrix-targeted treatments (≤ 0.6% of clusters in Phenotype 3 across other treatments; 8.9 – 12.9% of clusters in Phenotype 4 across other treatments). Protein-targeted biofilm matrix disruption releases the fewest large, oblate clusters (Phenotype 3, 0.3%) as compared to the other biofilm matrix-targeted treatments. Despite DNA-targeted biofilm matrix disruption releasing a higher relative concentration of cells (Fig. 2A), the distribution of cell clusters across the five phenotypes after DNase I treatment is similar to the distribution for the untreated condition (Table 2). EPS solubilization releases more mid-sized clusters (Phenotypes 2 and 5, 22.9% and 12.8%, respectively) and prolate clusters (Phenotype 5, 12.8%) than the other biofilm matrix-targeted treatments (Table 2). *S. epidermidis* biofilm matrix-targeted disruption therefore results in the release of cellular clusters with distinct size and morphology distributions across the five biofilm-detached cell cluster phenotypes.

**Table 2.**
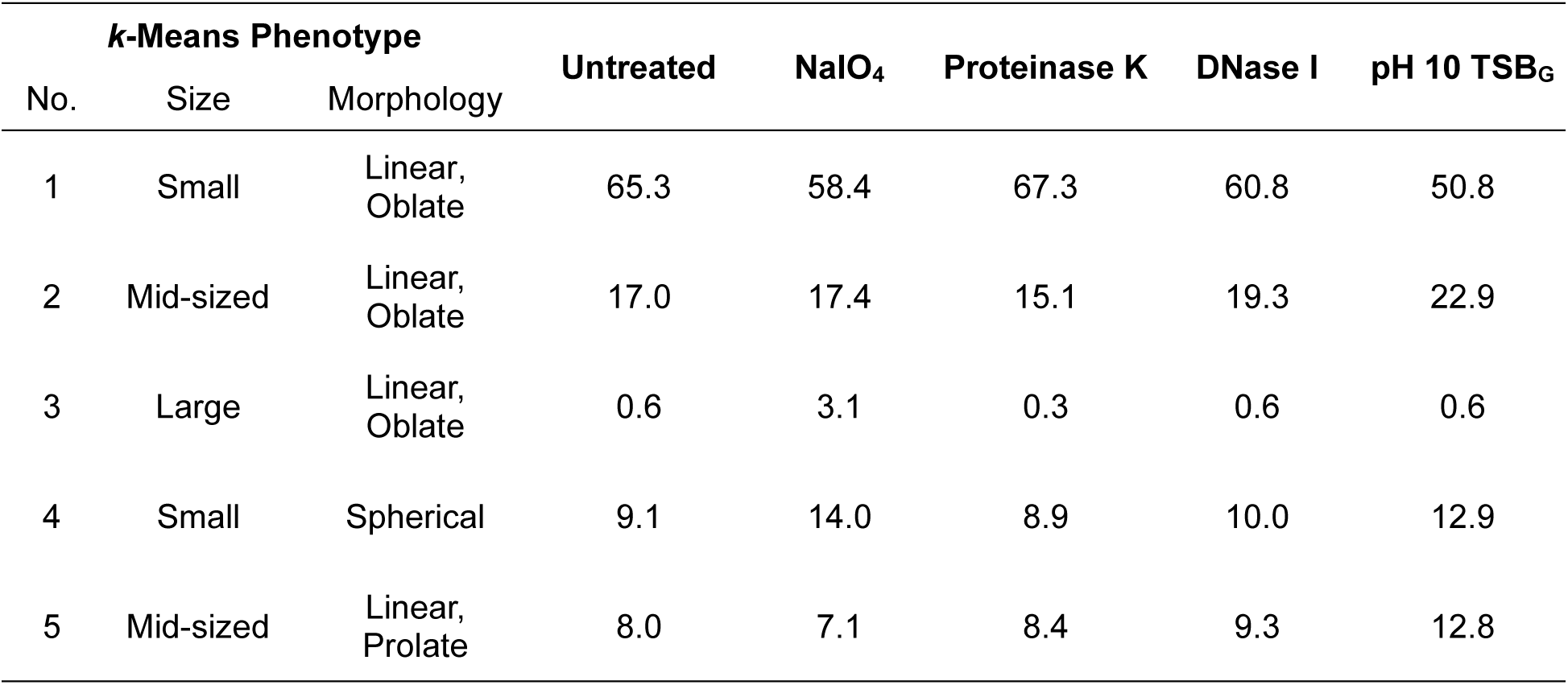
Percentages of *S. epidermidis RP62A* biofilm detached cell clusters within each biofilm-detached cell cluster phenotype across biofilm matrix-targeted treatments.

The relative abundance of cells released from *S. epidermidis* biofilms after matrix-targeted disruption varies with growth environment and between strains. The relative abundance of *S. epidermidis* RP62A cells detached from biofilms grown in flow cells and 96-well plates after PIA-targeted disruption and EPS solubilization are qualitatively similar; however, eDNA targeted disruption resulted in more detached cells for biofilms grown in flow cells than biofilms grown in 96-well plates (Fig. 4A). The relative abundance of detached cells after matrix-targeted biofilm disruption of *S. epidermidis* RP62A and three clinical isolates (P18, P37, P47) have qualitatively similar trends with a significantly higher abundance of cells released after PIA disruption and EPS solubilization compared to untreated biofilms (Fig. 4B). The relative abundance of detached cells after matrix-targeted treatment for four additional *S. epidermidis* strains (1457, P6, P2, P12) differ significantly from *S. epidermidis* RP62A (Fig. 4B) with PIA disruption only increasing the abundance of biofilm-detached cells for *S. epidermidis* P2. Thus, *S. epidermidis* biofilm growth environment and strain are factors that influence the release of biofilm cells due to matrix-targeted disruption.

**Figure 4.**
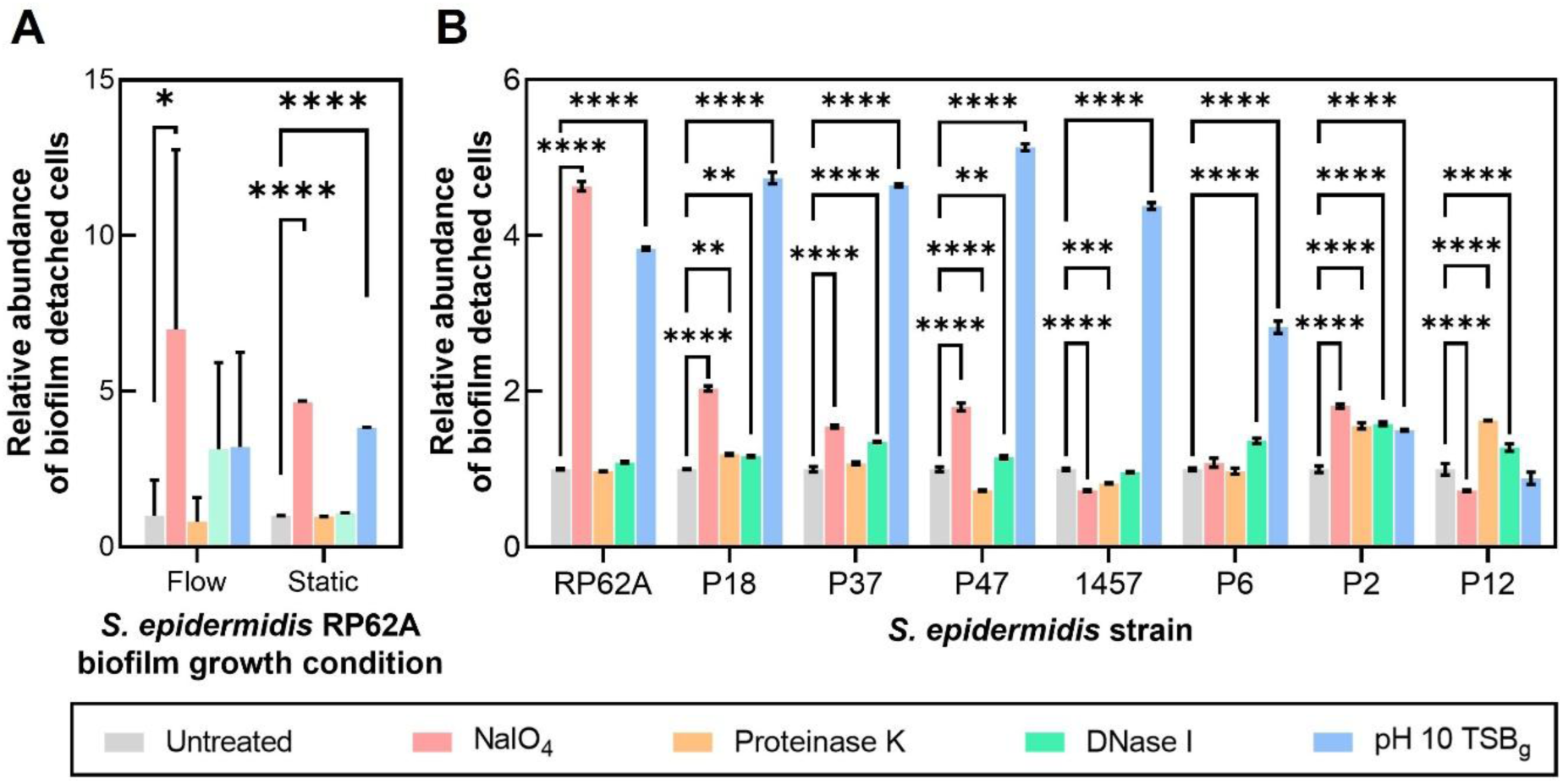
Relative abundance of *S. epidermidis* biofilm detached cells after matrix-targeted biofilm disruption across different growth conditions and strains. Relative abundance of biofilm detached cells, where the concentration or absorbance (OD_600_) of cells detached from biofilms after treatment is normalized to the untreated condition, for: (A) *S. epidermidis* RP62A biofilms grown in flow at a shear stress of 0.1 Pa or statically in a 96-well plate, and (B) eight *S. epidermidis* strains: RP62A, 1457, P2, P6, P12, P18, P37 and P47 grown in a 96-well plate. Statistical significance is determined by two-way ANOVA with Dunnett’s T3 multiple comparison tests (*P ≤ 0.05, **P ≤ 0.01, ***P ≤ 0.001, ****P ≤ 0.0001).

## Discussion

This work utilized quantitative image analysis of high-resolution confocal laser scanning microscopy images common to colloidal science (Fig. 1) to reveal how matrix-targeted disruption affects the concentration (Fig. 2A), size (Fig. 2B-C), morphology (Fig. 2D-E), and vancomycin susceptibility (Table S1) of cells and cellular clusters released from *S. epidermidis* biofilms. We also identify five distinct phenotypes of cells and cell clusters detached from *S. epidermidis* biofilms (Fig. 3, Table 1) by analyzing their biophysical properties with *k-*means clustering, an unsupervised machine learning algorithm. The predominant *S. epidermidis* biofilm-detached cell phenotype consists of small, linear, oblate clusters (1.9 ± 0.4 µm in diameter, 2-8 cells per cluster). All five *S. epidermidis* biofilm-detached cell cluster phenotypes are represented within the population of clusters released from all treatment conditions; however, the proportion of biofilm cell clusters from each phenotype varies uniquely with each biofilm matrix-targeted disruptor (Table 2). Additionally, the abundance of bacterial cells released from biofilms in response to matrix-targeted treatments varies with *S. epidermidis* biofilm growth environment and strain.

Five distinct *S. epidermidis* biofilm detached cell cluster phenotypes emerge from a heterogeneous dataset of 9,147 cell clusters using *k*-means clustering analysis with phenotypes that are stable and consistent across partitions of the data. Unsupervised machine learning techniques, including *k-*means clustering, are increasingly useful for uncovering hidden trends in heterogeneous biological datasets^42, 43^. In addition, the comparison of feature consistency across sub-samples of the dataset in a manner similar to *k*-fold cross validation^38^ is useful for evaluating model performance when analyzing heterogenous, experimental datasets. A similar unsupervised machine learning analysis approach could be applied to uncovering hidden trends of other inherently heterogenous characteristics of biofilms (e.g. biofilm cellular physiology^44^, structure^45^, or mechanics^46^).

The size and morphology of *S. epidermidis* biofilm detached cell clusters are important factors in the translocation and adhesion of biofilm-derived bacteria. Here we find that 79.8% of *S. epidermidis* RP62A biofilm-detached cells clusters are oblate and either small, mid-sized or large with respective average diameters of 1.9 ± 0.4 µm, 3.4 ± 0.8 µm, and 6.9 ± 2.1 µm. In addition, smaller subsets of bacterial clusters are detached as small, spherical (11.3%) and mid-sized, prolate clusters (8.9%) with diameters of 0.7 ± 0.1 µm and 2.6 ± 0.7 µm, respectively. Prior computational^47, 48^ and experimental studies^48^ have shown that 0.5-4 µm oblate particles are 10 to 100 times more likely to marginate and adhere to bloodstream walls across physiologically relevant shear stresses (0.5-6 Pa) than spherical or prolate particles with similar sizes^47, 48^. Additionally, larger particles (2-10 µm) marginate more rapidly to the bloodstream wall than smaller particles (∼500 nm) with similar shapes^49, 50^. Thus, bacterial clusters detached from *S. epidermidis* biofilms are released in morphologies and sizes that diversify cellular translocation by either promoting (Phenotypes 1-3, small, mid-sized, and large oblate clusters) or delaying (Phenotypes 4-5, small, spherical and mid-sized prolate clusters) the movement and adhesion of cell clusters to blood vessel walls. The size variability of the oblate phenotypes will also modulate the rate of biofilm detached cell cluster margination and adhesion to blood vessel walls. Because *S. epidermidis* is a non-motile organism^26^, the potential for size and morphology-based variations in the fluid dynamics of each biofilm detached cell cluster phenotypes is particularly impactful to understanding its translocation^51^. The current work is limited to studying physical properties of bacterial clusters; *S. epidermidis* surface adhesion molecules or the composition of remaining biofilm matrix in biofilm-detached cell clusters could be studied to reveal specific irreversible adhesion mechanisms or host cell interactions of biofilm detached cell clusters.

Size and morphology of *S. epidermidis* biofilm-detached cell clusters are also important to the host-immune response to bacteria. Successful phagocytosis by host immune cells varies with the size and shape of the target. The large, oblate *S. epidermidis* phenotype contains cell clusters with diameters ranging between 4.1 to 20.2 µm. The efficacy of polymorphonuclear leukocyte (PMN) phagocytosis decreases with increasing *S. epidermidis* cluster size, where 5, 10 and 15 µm cellular aggregates are successfully phagocytosed 89%, 40% and 5% of the time, respectively^18^. Thus, biofilm detached cell cluster size alone is likely to limit phagocytosis for a subset of the large, oblate clusters (Phenotype 3) whereas the small oblate and small spherical clusters (Phenotypes 1 and 4) have a high likelihood of being internalized by phagocytes. In addition, target shape, the angle of phagocyte approach, and target volume relative to phagocyte size are all important parameters for successful phagocytosis^19^. Macrophage phagocytosis of ellipsoidal particles with volumes up to the size of the macrophage itself is typically successful when macrophage approach is along the minor axis of an ellipsoid^19^; however, when macrophage approach is along the major axis of both oblate or prolate ellipsoids, the macrophage typically fails to internalize the particle and “frustrated” phagocytosis occurs even for particles that are only 0.2% of the macrophage volume^19^. Thus, when phagocyte approach is along the major axis of cell clusters from the mid-sized oblate, large oblate, and mid-sized prolate biofilm-detached cell phenotypes (Phenotypes 2, 3, 5), “frustrated” phagocytosis is likely to occur. While the current work is limited to size and morphology of cell clusters, prior work has shown that PMNs can successfully engulf fragments of large *S. epidermidis* aggregates^18^, suggesting an interplay between bacterial cluster size, morphology and mechanics on phagocytosis success. Biofilm-detached cell cluster mechanics could be evaluated in future work to further advance the understanding of the host immune cell response to biofilm-detached cell clusters.

All five *S. epidermidis* biofilm detached cell cluster phenotypes are present after the four matrix-targeted disruption strategies studied here; however, differences in the proportions of each phenotype indicate that there are distinct biofilm matrix interactions that contribute to how biofilm cells and matrix materials assemble and disassemble. Polysaccharide-targeted *S. epidermidis* biofilm disruption contains the highest proportion of large, oblate (3.1%) and small, spherical (14.0%) bacterial clusters, indicating that PIA disruption results in the sloughing of larger cell clusters and erosion of more individual cells than in untreated biofilms. Protein-targeted *S. epidermidis* biofilm disruption results in a modest decrease in mid-sized, oblate clusters (17.0% to 15.1%) and large, oblate clusters (0.6 % to 0.3%). This suggests that matrix protein interactions within the biofilm milieu play an important role in achieving large- and mid-sized, oblate cell clusters detached from biofilms, potentially through protein-mediated cell-EPS adhesion mechanisms^5^. Removal of DNA from the matrix results in increased quantities of mid-sized clusters and increased erosion of individual biofilm cells compared to untreated biofilms—as evidenced by modest increases in the proportion of cell clusters released as mid-sized oblate (17.0% to 19.3%), mid-sized prolate (8.0% to 9.3%), and small spherical bacterial clusters (9.1% to 10.0%). After increasing *S. epidermidis* biofilm pH to induce EPS solubilization, there is a higher portion of mid-sized oblate (17.0% to 22.9%), mid-sized prolate (8.0% to 12.8%), and smaller spherical clusters (9.1% to 12.9%) detached from biofilms; this trend is similar to that of the DNA-targeted biofilm disruption, but with larger increases in the proportions of cells in each phenotype. This result suggests that eDNA plays a significant role in modulating the previously reported pH-induced phase stability of the *S. epidermidis* biofilm matrix, where the matrix is more stable (soluble) in solution with increasing pH^9^. This advanced understanding of *S. epidermidis* biofilm disassembly in response to matrix-targeted treatments can be applied to develop mechanistic models of biofilm development and disassembly, advance the understanding of how biofilm infections develop into systemic infections, and predict the efficacy of biofilm removal and treatment strategies.

Finally, the abundance of *S. epidermidis* biofilm cells released after matrix-disruption can differ with growth environment and between strains. *S. epidermidis* RP62A biofilms grown at a shear stress of 0.1 Pa release more cells after PIA-targeted disruption and DNA-targeted disruption than biofilms grown statically (Fig. 4A) suggesting higher variability of PIA concentrations and higher quantities of eDNA in the matrix of *S. epidermidis* biofilms grown at 0.1 Pa. *S. epidermidis* RP62A is known to increase PIA production with increasing shear stress^52^; thus, our finding is consistent with prior results. Across 50% (4/8) of *S. epidermidis* strains in this study, PIA-targeted treatment and pH-induced matrix solubilization has a significant impact on the abundance of cells released from biofilms, indicating similarities across strains in efficacy of biofilm disruption strategies. As 20-60% of *S. epidermidis* clinical isolates carry the *ica* operon responsible for PIA production^53^, a 50% success rate for PIA-targeted biofilm disruption is unsurprising. On the other hand, PIA-targeted treatment of *S. epidermidis* 1457 did not significantly modulate the abundance of released biofilm cells, which was unexpected as *S. epidermidis* 1457 typically forms PIA-rich biofilms^54^. The efficacy of matrix-targeted biofilm disruptors is variable for the remaining three *S. epidermidis* strains studied. The abundance of cells detached from *S. epidermidis* P12 biofilms is significantly increased with Proteinase K and DNase I treatments; *S. epidermidis* P6 biofilms are significantly disrupted by only DNase I treatment; and all biofilm-disruption treatments significantly increase the abundance of cells released from *S. epidermidis* P2 biofilms (Fig. 5). *S. epidermidis* biofilm development can be mediated by multiple molecules, including PIA, proteins, and DNA, where expression of these molecules can vary with strain and growth environment^55^. As shown here, *S. epidermidis* biofilm-matrix targeted disruptor efficacy varied across two growth environments and eight strains. Thus, variations in *S. epidermidis* biofilm matrix composition based on growth environment and strain should be considered when evaluating properties of biofilm-detached cell clusters and the efficacy of biofilm matrix-targeted disruption techniques.

## Conclusions

Our work determined that *S. epidermidis* cells and cell clusters released from untreated biofilms and biofilms treated with matrix-targeted disruptors have five distinct phenotypes with the largest fraction of cell clusters (60.4%) in the small, linear, oblate phenotype, where clusters are 1.9 ± 0.4 µm in diameter and contain 2-8 cells per cluster. Five previously unknown phenotypes were revealed using *k-*means clustering, an unsupervised machine learning technique for uncovering hidden trends in data. The distribution of *S. epidermidis* biofilm detached cell clusters across the five phenotypes varied with matrix-targeted disruption technique. Trends in the abundance of *S. epidermidis* cells released from biofilms after matrix targeted disruption also varied with growth condition and strain. Advancing the understanding of the physical properties of *S. epidermidis* cell clusters detached from biofilms is an important step in understanding bacterial survivability, translocation, and susceptibility to the host immune response as well as in the development of matrix-targeted biofilm control strategies.

## Supplementary Information

Supplementary information is provided as a separate file online. The supplementary information contains Supplementary Figures S1-S9 and Supplementary Table S1.

## Acknowledgements

We thank J. S. VanEpps (Departments of Emergency Medicine and Macromolecular Science and Engineering, University of Michigan) for providing *S. epidermidis* P2, P6, P12, P18, P37 and P47.

## Contributions

List of contributions using CRediT taxonomy.

**S.R.P** – Conceptualization, data curation, formal analysis, investigation, methodology, software, validation, visualization, writing – original draft, writing – review and editing

**B.P.** – Formal analysis, methodology, writing – review and editing

**G.B.** – Investigation, writing – review and editing

**R.P** – Formal analysis, methodology, validation, writing – review and editing

**E.J.S.** – Conceptualization, formal analysis, funding acquisition, methodology, project administration, supervision, validation, visualization, writing – original draft, writing – review and editing.

## Declaration of generative AI and AI-assisted technologies in the manuscript preparation process

During the preparation of this work the authors used ChatGPT 4.0 to assist in the development and optimization of Python scripts for image analysis. After using this tool, the authors reviewed and edited the generated code as needed and take full responsibility for the content of the published article.

