## Supplementary Information for "Biophysical properties and phenotypes of cell clusters detached from *Staphylococcus epidermidis* biofilms after matrix-targeted disruption"

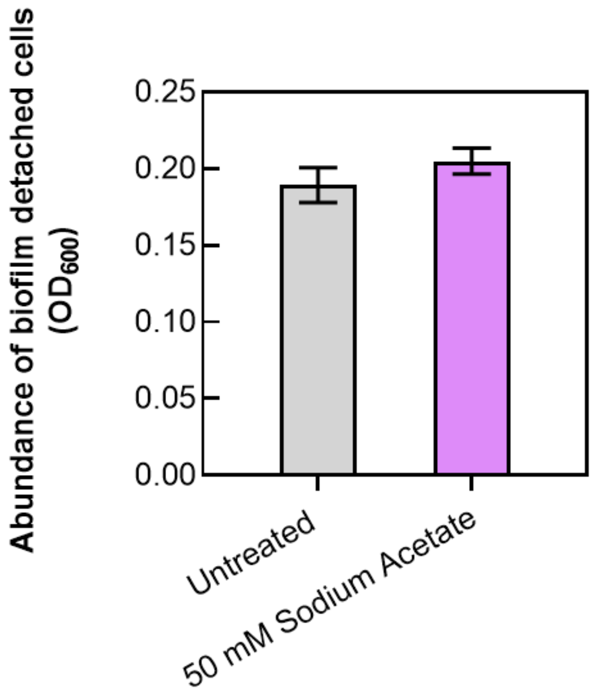


**Figure S1. Evaluation of the effect of 50 mM sodium acetate buffer on the abundance of cells detached from *S. epidermidis* RP62A biofilms.** Application of 50 mM sodium acetate to biofilms for 1 hour does not significantly impact the abundance of cells detached from *S. epidermidis* RP62A biofilms compared to untreated biofilms ($P = 0.33$, Welch's t-test). Abundance of detached cells is measured as $OD_{600}$. The polysaccharide-targeted matrix disruptor, 10 mM sodium m-periodate ($NaIO_4$), is suspended in a 50 mM sodium acetate oxidation buffer; thus, the effect of 50 mM sodium acetate oxidation buffer on *S. epidermidis* biofilms was evaluated independent of $NaIO_4$. $NaIO_4$ is suspended in a 50 mM sodium acetate oxidation buffer instead of $TSB_G$ as $NaIO_4$ can target the glucose in $TSB_G$ rather than PIA in the biofilm matrix[1] when glucose is present in the growth media.

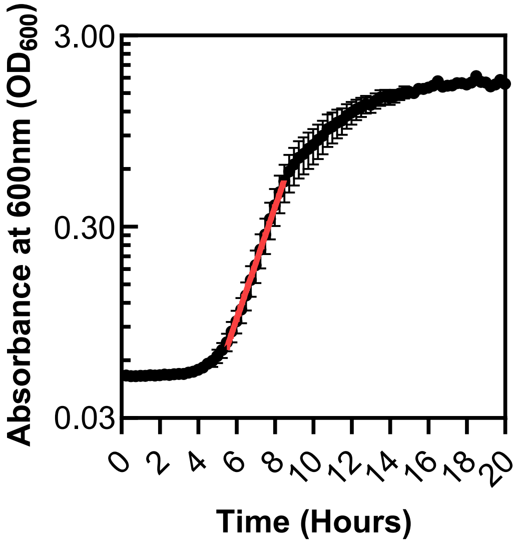


**Figure S2. Determination of doubling time of *Staphylococcus epidermidis* RP62A.** Growth curve of planktonic *S. epidermidis* RP62A in $TSB_G$ at 37°C, 230 RPM. Doubling time was calculated to be 62 minutes with a 95% confidence interval using best fit for an exponential (Malthusian) growth model in GraphPad Prism 9.3.1.

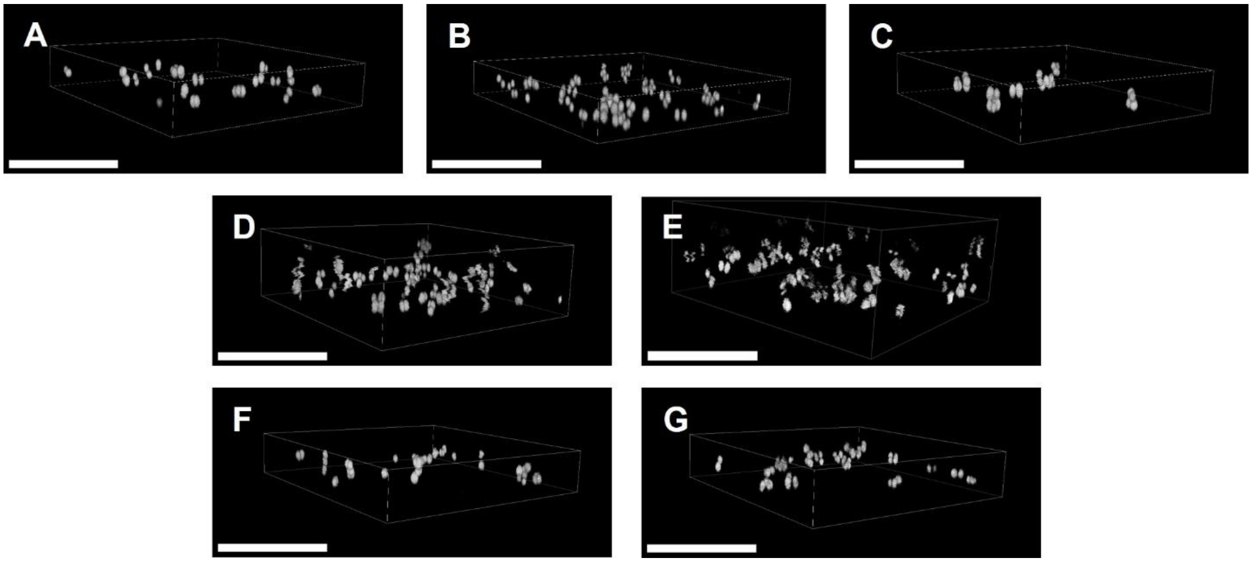


**Figure S3. Imaging of biofilm detached bacterial cell clusters after biofilm matrix-targeted treatments.** Cell clusters detached from (A) untreated biofilms and biofilms treated with (B) $NaIO_4$ or (C) Proteinase K were immobile at the coverslip after 1 hour (30 minutes staining and 30 minutes of settling time) on untreated coverglass. *S. epidermidis* RP62A cell clusters detached from biofilms treated with (D) DNase I or (E) pH 10 $TSB_G$ remained mobile at the coverslip after 1 hour on untreated coverglass as indicated by noise in 3D renderings. (F) *S. epidermidis* RP62A cell clusters detached from biofilms treated with DNase I were immobile after 1 hour on coverglass treated with oxygen plasma (100 watts, 45 seconds), a surface treatment that increases the coverglass surface potential. (G) *S. epidermidis* RP62A cell clusters detached from biofilms treated with pH 10 $TSB_G$ were immobile after 1 hour on a coverglass treated with Rain-X Glass Water Repellent (Illinois Tool Works, Inc., USA), a hydrophobic surface treatment. (SBs = 20 μm).

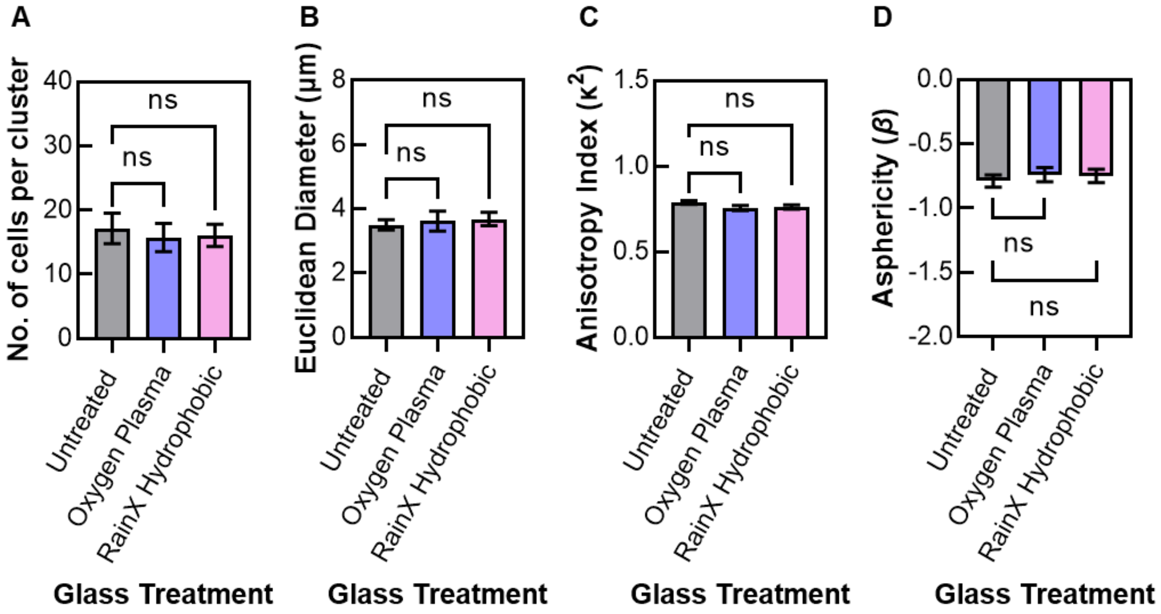


**Figure S4. Physical properties of planktonic *S. epidermidis* RP62A cell clusters are not significantly altered by coverglass treatments.** (A) Number of cells per cluster, (B) Euclidean diameter, (C) anisotropy index and (D) asphericity of *S. epidermidis* RP62A cell clusters from mid-log growth planktonic cell culture ($OD_{600}$ = 0.06) in untreated, oxygen plasma treated, or Rain-X Glass Water Repellent treated Nunc Lab-Tek 8-Well Chambered Coverglass dish (Thermo Scientific™ 155409). No significant differences were observed between number of cells per cluster, Euclidean diameter, anisotropy index and asphericity of planktonic cell clusters on untreated, oxygen plasma treated and Rain-X treated coverglass ($P > 0.05$, non-parametric Kruskal-Wallis with Dunn's multiple comparison tests; 'ns' denotes no significance).

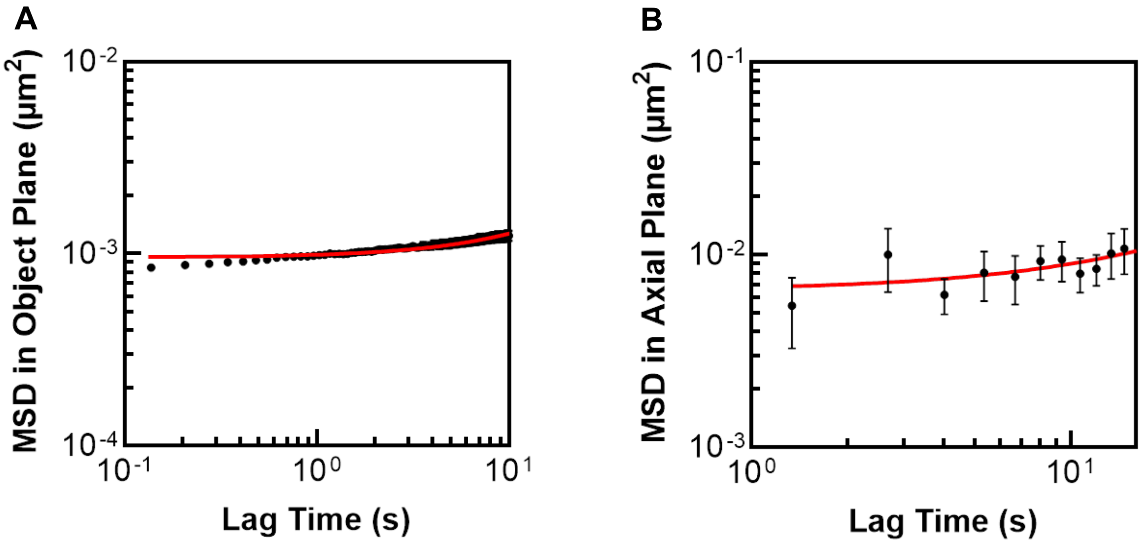


**Figure S5. Static error for 3D particle identification in CLSM image volumes.** The static error, $\sigma$, is represented as a constant offset in the measured mean squared displacement (MSD) for immobile particles, where $MSD_{measured,\ t=0} = 2\sigma^2$. The MSDs of immobilized 1.0 μm diameter polystyrene microspheres (Polysciences, Warrington, PA, USA) were determined and used to compute the static error in the object and axial planes using the methods of Savin and Doyle[2]. Briefly, the polystyrene beads were arrested at the coverslip of 8-well Nunc Lab-Tek II chambered coverglass dishes via immersion in 100 mM sodium chloride. Next, five xy-t time series images collected at 14.54 frames per second were used to evaluate (A) the MSD of the arrested polystyrene particles in the object plane (xy). Five xyz-t time series images at 0.746 frames per second were analyzed to evaluate (B) the MSD of the arrested polystyrene particles in the axial plane (xz). The line of best fit for the MSD is $MSD_{measured} = 2.0 \times 10^{-5} \cdot t + 1.0 \times 10^{-3}$ in the object plane, and $MSD_{measured} = 1.9 \times 10^{-4} \cdot t + 6.2 \times 10^{-3}$ in the axial plane in units of μm$^2$. At t = 0, $MSD_{measured,\ t=0}$ is $1.0 \times 10^{-3}$ μm$^2$ in the object plane and $6.2 \times 10^{-3}$ μm$^2$ in the axial plane. Thus, the static error in particle location was determined to be ± 23 nm in the object plane (xy) and ± 56 nm in the axial plane (xz).

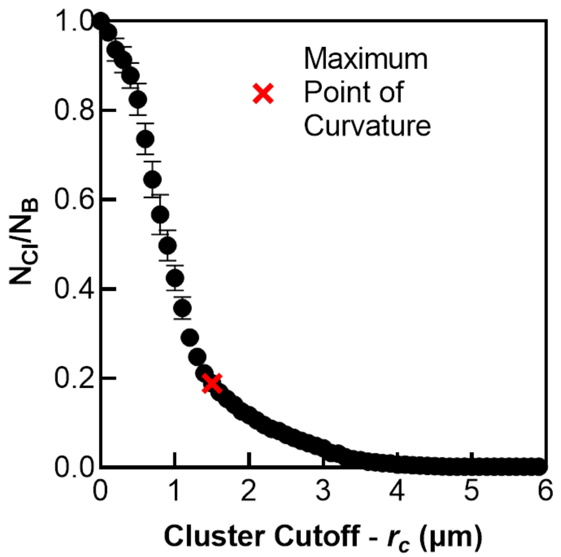


**Figure S6. Determination of cluster cutoff distance for grouping biofilm detached cells into bacterial cell clusters.** The optimal cluster cutoff distance, $r_{c,optimal}$ was determined to be 1.5 µm based on the inflection point, or rather the maximum point of curvature (red x), on the cluster cutoff distribution curve. The cluster cutoff distribution curve is a plot of the ratio of the number of bacterial clusters ($N_{Cl}$) to the total number of bacterial cells ($N_B$) within the analyzed image volumes against cluster cutoff distance ($r_c$). The inflection point occurs where the rate of change in the ratio of $N_{cl}/N_B$ is at a minimum value[3].

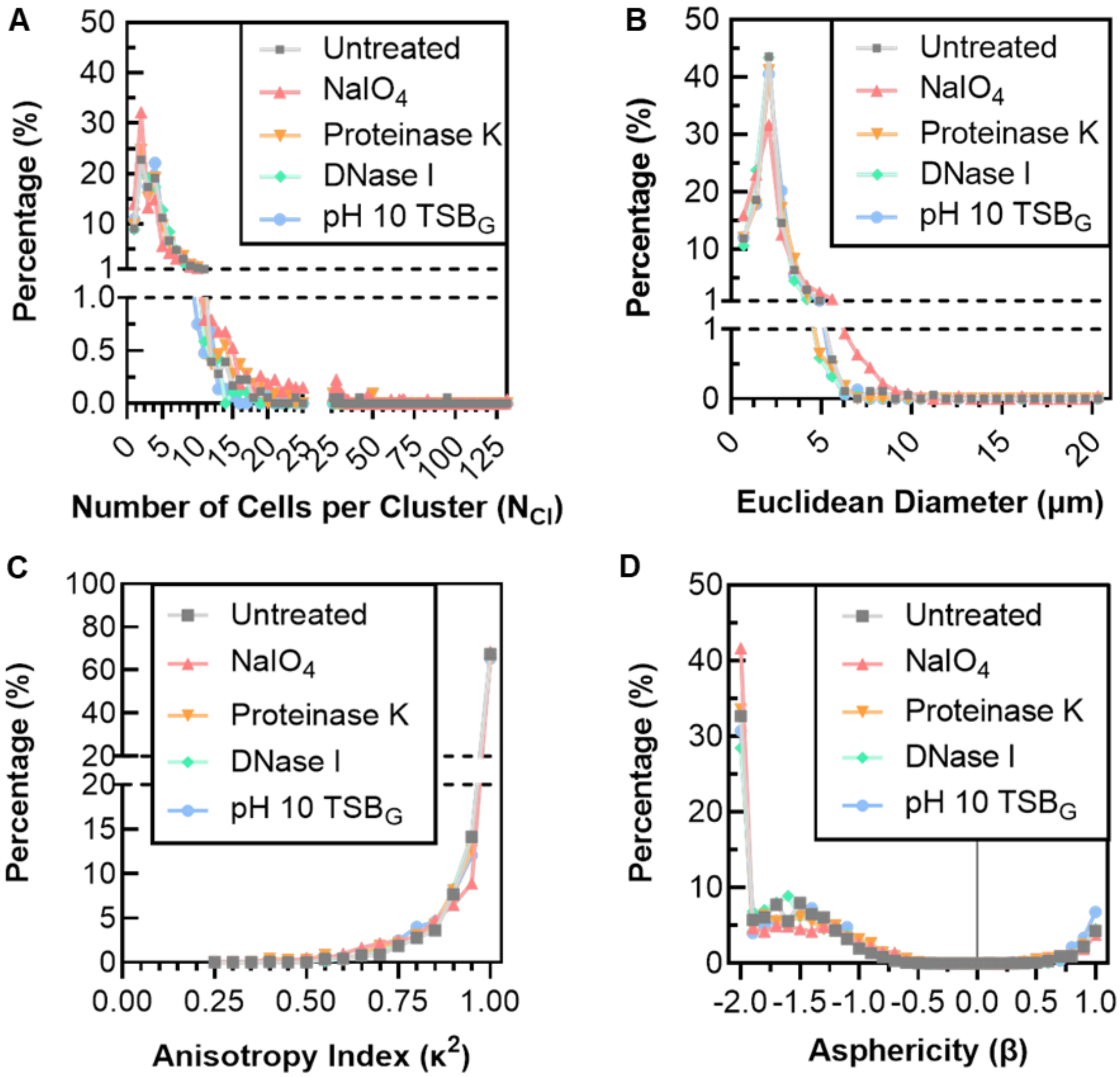


**Figure S7. Histograms of physical properties of bacterial clusters detached from *Staphylococcus epidermidis* biofilms after matrix-targeted disruption are broad and non-Gaussian.** Histograms of (A) number of cells per cluster, (B) Euclidean diameters, (C) anisotropy indices and (D) asphericities of cell clusters detached from *S. epidermidis* RP62A biofilms after no treatment or treatment with $NaIO_4$, Proteinase K, DNase I or pH 10 $TSB_G$. All distributions deviated significantly from normality, as confirmed by Shapiro-Wilk tests (all $P < 0.05$) and visual inspection of histograms.

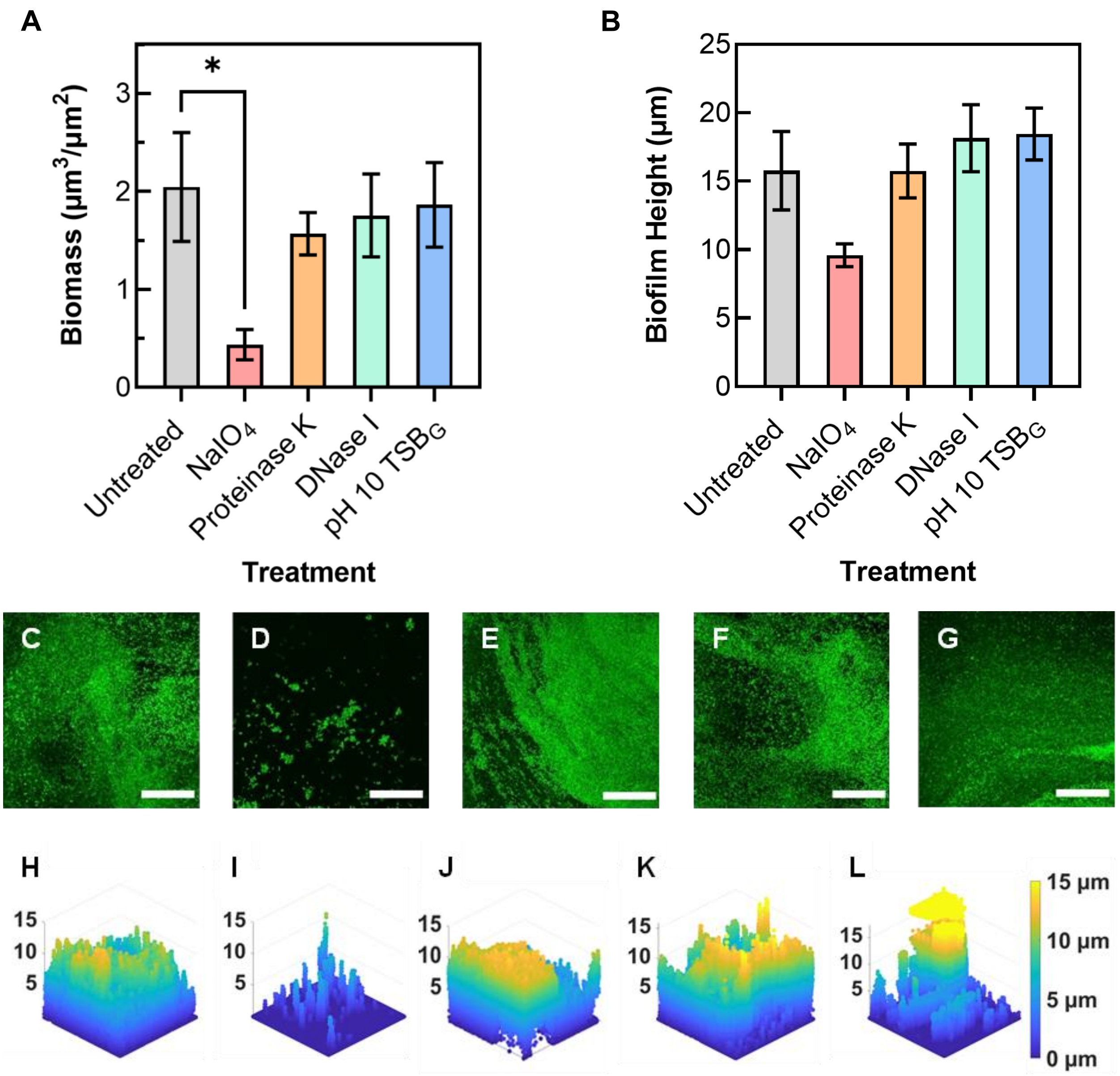


**Figure S8. Effect of matrix-targeted disruptors on *S. epidermidis* RP62A biofilm remaining in flow cell after treatment.** *S. epidermidis* RP62A (A) biofilm biomass and (B) biofilm height after no treatment or treatment with $NaIO_4$, Proteinase K, DNase I, or pH 10 $TSB_G$. Statistical significance is determined by one-way ANOVA with Dunnett's T3 multiple comparison tests (*P < 0.05). Representative (C-G) maximum intensity projections and (H-L) 3D renderings of *S. epidermidis* RP62A biofilm remaining in flow cells after: (C,H) no treatment, or treatment with (D,I) $NaIO_4$, (E,J) Proteinase K, (F,K) DNase I or (G,L) pH 10 $TSB_G$. SBs = 50 μm. 3D renderings are from CLSM image volumes with dimensions of 184.88μm x 184.88μm x 15μm.

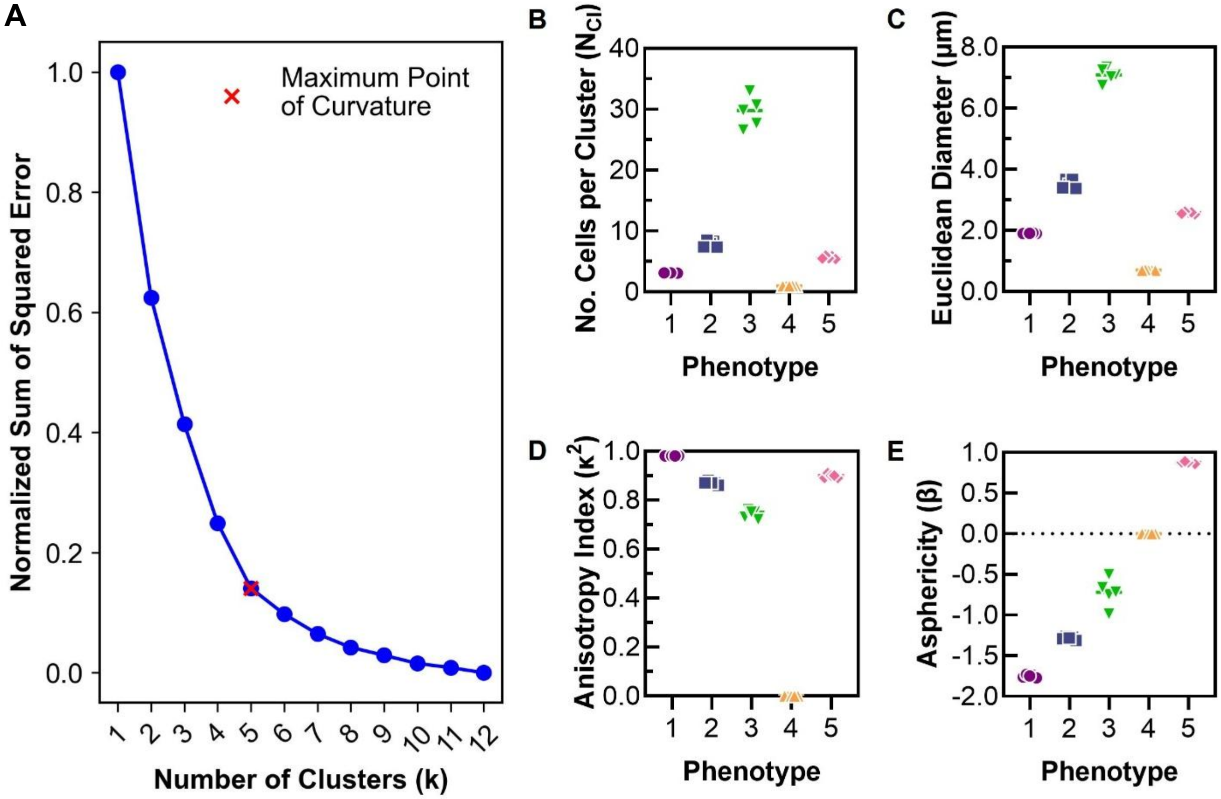


**Figure S9. Determination of *k*-value for *k*-means clustering analysis of biofilm-detached cell clusters.** (A) Plot of *k*-value against the normalized sum of squared error. The optimal *k*-value was identified as five using the Elbow method, a common clustering algorithm heuristic that identifies an optimal *k*-value by evaluating the maximum point of curvature (red x) on a plot of the Sum of Square Error (SSE) against a range of *k*-values[4]. (C-F) The average (C) number of cells per cluster, (D) Euclidean diameter, (E) anisotropy index, and (F) asphericity of the five *S. epidermidis* RP62A biofilm-detached cell cluster phenotypes are consistent across five sub-samples of the dataset using a *k*-value of 5. Each point is a sub-section of data.

**Table S1.** Vancomycin minimum inhibitory concentration (MIC) of cells released from *S. epidermidis* RP62A biofilms after biofilm matrix-targeted disruption.

| Matrix disruption agent | Vancomycin MIC (µg/mL) |
|---|---|
| Untreated | 2.0 |
| $NaIO_4$ * | --- |
| Proteinase K | 2.0 |
| DNase I | 2.0 |
| pH 10 $TSB_G$ | 2.0 |
| Planktonic cells | 2.0 |

*Note: Bacterial cells detached from biofilms were not viable after treatment with NaIO4.
